# A High-Throughput Sampling Method for Detection of *Meloidogyne enterolobii* and Other Root-Knot Nematodes in Sweetpotato Storage Roots

**DOI:** 10.1101/2023.05.10.540019

**Authors:** Julianna Culbreath, Catherine Wram, Churamani Khanal, Tyler Bechtel, Phillip A. Wadl, John Mueller, William B. Rutter

## Abstract

*Meloidogyne enterolobii* is an aggressive root-knot nematode (RKN) species that has emerged as a significant pathogen of sweetpotato in the Southeastern United States. *M. enterolobii* is spread primarily through the movement of infected ‘seed’ sweetpotatoes used for propagation. The RKN resistance in commercially grown sweetpotato cultivars has proven ineffective against this nematode. Detecting RKN in sweetpotato by eye is unreliable, and further distinguishing *M. enterolobii* from other RKN species that infect sweetpotato is labor intensive; relying on molecular tests conducted on individual nematodes dissected out of host roots by trained technicians. Here, we have developed a high-throughput survey method to collect skin samples and extract total DNA from batches of sweetpotato storage roots. Combining this method with species-specific PCR assays allowed for quick and sensitive detection of *M. enterolobii* and other RKN species infecting sweetpotatoes. We tested this method using batches of infected storage roots at varying levels of *M. enterolobii* infection. We also inoculated skin samples with varying numbers of individual *M. enterolobii* eggs to determine the method’s detection threshold and used this method to conduct surveys for RKN on fresh market sweetpotatoes. Our results show that this method can consistently and reliably detect *M. enterolobii* in sweetpotato batches at levels as low as 2 eggs per 10 mL skin sample. This method will be a useful tool to help screen for the presence of *M. enterolobii* in ‘seed’ sweetpotatoes before they are replanted, thereby helping to slow the spread of this nematode to *M. enterolobii*-free sweetpotato growing operations.

## 1. Introduction

Sweetpotato (*Ipomea batatas*) is a staple root crop in subtropical and tropical regions around the world. In 2021, the United States (US) sweetpotato industry produced 1.6 million US tons of storage roots with an estimated value of $600 million dollars, which is nearly double the1 production volume and triple the profit compared to 2000 (NASS 2021). Sweetpotato is a clonally propagated crop, which requires planting ‘seed’ storage roots in the early spring for shoot production. Shoots are cut into sections termed ‘slips’, which are used to plant production fields. The movement of sweetpotato seed roots and slips can vector the spread of soilborne plant pathogens, including root-knot nematodes (RKN, *Meloidogyne* spp.) (Overstreet et al. 2019; Quesada-Ocampo 2018, 2019; Silva et al. 2021).

RKN are sedentary endoparasites that infect a wide range of crops, and cause millions of dollars in yield losses each year (Perry and Moens 2009). These pests migrate into the roots of a host plant and establish elaborate feeding sites within the root (Chitwood 1949). After establishing its feeding site, the nematode will remain inside the root for the rest of its life cycle (Krusberg L. R. and Nielsen 1958; Stirling et al. 2020). The juveniles develop into mature females which can each produce hundreds of eggs (Karssen et al. 2013). In addition to acting as a nutrient sink, RKN can also induce galling and cracking in the surrounding root tissue. These disruptions on the surface of the crop can ultimately render root vegetables, like sweetpotato, unmarketable (Lawrence et al. 1986; Osunlola and Fawole 2015). A single RKN-infected sweetpotato root planted in a field can initiate an infestation that will eventually spread over the entire cropping area. Once RKN is established in a field, it is nearly impossible to eradicate from that field (Baidoo et al. 2017; Clark et al. 2013; Stirling et al. 2020). The wide host range of RKN includes many row crops and weed species which contributes to the spread and survival of RKN in a field.

In recent years, a particularly aggressive species of RKN, *M. enterolobii,* has spread across sweetpotato production operations in the Southeastern US. *M. enterolobii* was first described on pacara plants in China in 1983 (Yang & Eisenback 1983). It was first reported in Florida in 2002, and has since spread to North Carolina, South Carolina, and Georgia (Ye et al. 2013, 2018; Rutter et al. 2019; N.C. Department of Agriculture and Consumer Services 2017; Brito et al. 2004). Instances of *M. enterolobii* introduced into sweetpotato fields by infected seed roots have recently been reported in Louisiana, Florida, and Georgia, and the nematode has been reported in all sweetpotato growing counties in North Carolina (FINDMe 2022; Gu et al. 2021; Hare 2019; Rezende et al. 2022). *M. enterolobii* has a broad host range apart from sweetpotato and is known to infect and damage the majority of crops grown in the Southeastern US (Philbrick et al. 2020; Castagnone-Sereno 2012). Moreover, *M. enterolobii* can successfully reproduce on resistant crop varieties that have historically been used to manage endemic species such as *M. incognita* (Kiewnick et al. 2009; Silva et al. 2019; Rutter et al. 2021). Reducing the movement of *M. enterolobii* infected storage roots would help mitigate the impact of this nematode on US agriculture.

Currently, there is no established method for certifying sweetpotato storage roots as free of RKN. Traditional methods used to detect and speciate RKN in root vegetables often involve visual inspection of roots and isolation of individual nematodes. These approaches are labor intensive, require specialized nematology training, and may not provide an accurate representation of all species present in a mixed population of RKN. In this study, we have developed and tested a new sampling method that can reliably detect the presence of *M. enterolobii* and other RKN species in batches of sweetpotato storage roots. The results indicate that this method could be a useful approach to quickly survey storage roots for the presence of *M. enterolobii* and potentially reduce the number of infected ‘seed’ potatoes planted in the field.

## 2. Materials and methods

### 2.1 Sweetpotato storage root material

Negative control (uninfected) sweetpotatoes were harvested from clean sweetpotato clones (‘Beauregard’, ‘Covington’, ‘Regal’, USDA-10-185, or ‘W-353’) which had been grown for 120 days in sterile soil within a controlled Plant Biosafety Level Two (BL2-P) greenhouse environment (approximately 12 hour photoperiod with temperatures ranging between 16.7° C – 32.1° C) at the USDA, ARS, US Vegetable Laboratory in Charleston, SC.

Positive control (*M. enterolobii-*infected) storage roots of either a *M. enterolobii* susceptible cultivar (Beauregard), resistant cultivar (Regal) or resistant breeding line (USDA 10-185) were used to assess the consistency and sensitivity of the sampling method (Rutter et al. 2021). Infected controls were produced by planting sweetpotato slips in sterile soil and allowing slips to grow for four weeks before inoculation with 10,000 *M. enterolobii* (isolate SC.1) eggs. The plants grew for 120 days under the conditions described above.

To validate the newly developed skinning method, these positive and negative control roots were processed in different batch combinations. Each batch was between 5-10 lbs and contained at least 5 storage roots. These batches were classified into four different test groups based on their known level of *M. enterolobii* infection: (1) heavily infected control batches of susceptible sweetpotatoes all showing obvious damage, (2) moderately infected control batches, consisting of a single heavily infected susceptible root added into a batch of asymptomatic roots from a local grocery store, (3) minimally infected control batches of resistant sweetpotatoes which showed little to no galling symptoms, and (4) uninfected control batches of sweetpotatoes used as negative controls.

For surveys, batches of up to 10 lbs of sweetpotato storage roots were collected from supermarkets, packing houses, or grower fields between May 2021 and October 2022. When available, the origin and producer information was recorded. Twenty-nine survey samples were collected and are described in Table 2. Each sample was stored in a leak-proof container at 4 °C until skinning.

### 2.2 Storage root skinning and sampling method

Each batch of storage roots was placed in an autoclave-sterilized 5-gallon plastic bucket (Home Depot, Model #05GLHD2), containing enough tap water to cover the roots. They were then skinned for 5 min using an autoclave-sterilized abrasive bristle brush (Ryobi, Model #A95GCK1) attached to a corded electric drill (RIDGID, Model #R71111) (Fig. 1a). Once the roots were skinned, the skinned roots were removed, and the mixture of skin material and water was poured through stacked stainless-steel 8” sieves (No. 270 over No. 500, Gilson Company Inc.). The sieves were placed over a second bucket and agitated with a sieve shaker (Gilson Company Inc., Model SS-23) to separate the water from the skinnings (Fig. 1b). Once the water had drained from the skinnings, the contents of the top and bottom sieves were mixed on the bottom (No. 500) sieve with a sterile spoon, and two 10 mL samples of the skinnings were transferred to separate sterile 50 mL conical bottom tubes (VWR, Catalog #76204-404). All skinning samples were stored at -80 °C until DNA extraction (Fig. 1c) (Supplemental Film 1, <AGDATA commons link to be added>).

**Figure 1.**
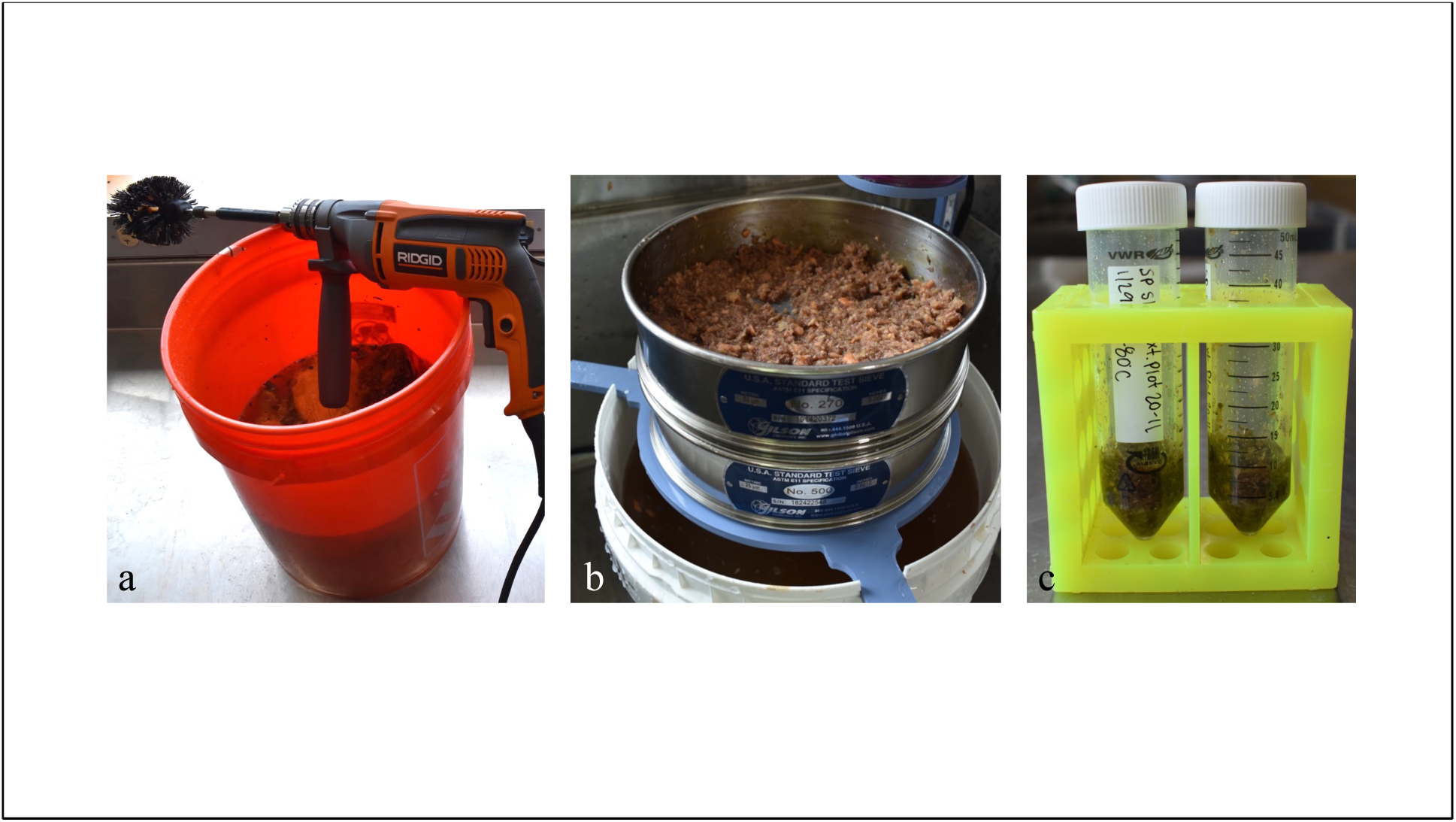
The equipment used to collect samples for high-throughput detection of *Meloidogyne* species in sweetpotato storage roots. (a) Abrasive bristle brush attached to a corded electric drill, peeling a 10 lb batch of sweetpotatoes in a 5-gal bucket with water. The liquid contents are then transferred to the collection sieves that retain the skinning’s for sampling and *post-hoc* egg extraction. (b) The combined skinnings collected in the sieve stacks from a 5-min peeling. (c) A 10mL sample collected from the mixed sweetpotato skins.

Total DNA was separately extracted from each 10 mL skinning sample using the DNeasy^®^ *mericon*^®^ Food Kit (Qiagen, Hilden, Germany). Each extraction was performed following the manufacturer’s instructions for standard large-scale extractions with minor modifications. Briefly, samples were first homogenized to a fine powder in liquid nitrogen using a mortar and pestle. DNA was extracted from 2 g of the fine powder from each ground sweetpotato skin sample and was eluted in a final volume of 100 μL elution buffer.

### 2.3 Diagnostic PCR and qPCR for skin samples

Quantitative real-time PCR (qPCR) was performed for all total skin DNA samples using *M. enterolobii* specific primers and a locked nucleic acid (LNA) probe as previously described (Kiewnick et. al 2015). A reaction volume of 20 µL was composed of 10 µL Roche LC480 Probes Master, 1 µL of LNA probe 17, 1 µL of Ment17F, 1 µL of Ment17R, 4 µL of template DNA, and 3 µL of nuclease-free water (see Supplemental Tables 1 & 2 for primer sequences and reaction conditions). All reactions were quantified on a Roche LC480 Light cycler instrument (LC480) (Roche, Basel, Switzerland). Three technical replicates were used for each sample.

Conventional PCR was also performed on the same total skin DNA samples using two sets of general RKN primers TRNAH/MRH106 and MORF/MTHIS as previously described (Baidoo et al. 2016, Pagan et al. 2015) (Supplemental Table 1). When the results of the general RKN primer sets indicated the presence of *M. enterolobii,* a second species specific primer set MeF/MeR was used to confirm the results (Hu et al. 2011). If an RKN species other than *M. enterolobii* was detected, no further molecular analysis was performed, and we classified it as positive *Meloidogyne spp*. A no template control (NTC) was included alongside each diagnostic PCR assay. Amplifications with primer sets were performed in a final volume of 25 μL using 12.5 μL 2X Apex *Taq* red master mix (Genesee Scientific, San Diego, CA), 1.25 μL each of 10-μM forward and reverse primers and 3 μL of DNA template. Reactions were performed using a Bio-Rad T100 thermal cycler, following primer specific conditions (Supplemental Table 2).

### 2.4 Detection sensitivity assays

Skinning samples were collected from batches of uninfected control roots as described above. *M. enterolobii* eggs were then inoculated directly into fresh, uninfected 10 mL skinning samples at different inoculation densities (0, 2, 10, or 100 eggs/10 mL) before freezing the entire sample at -80 °C. Three skinning samples (biological replicates) for each of the 4 inoculation densities were extracted for DNA as described above for a total of 12 samples. The eggs used for inoculation were freshly extracted from an infected greenhouse *M. enterolobii* (isolate SC.1) culture grown on the tomato cultivar Rutgers and counted under a dissecting microscope as previously described (Hussey and Barker 1973). We quantified the amount of *M. enterolobii* tissue in each inoculated sample using qPCR as in Section 2.3. The standard curve for eggs per 10 mL skin sample was generated using the lm function in R version 4.2.0. The results were plotted using the *ggplot2* package in RStudio version 2022.2.2.485 (RStudio Team 2022; Wickham 2016).

### 2.5 *Post-hoc* bioassays

*Post-hoc* bioassays were completed for 14 negative control batches, 10 positive control batches (4 moderately infected and 6 minimally infected), and all 29 survey batches. After collecting samples for DNA extraction of the sweetpotato skins, we used the leftover skinning material to perform *post-hoc* bioassays by extracting eggs using well established methods (Hussey and Barker 1973). Briefly, the leftover skinnings were placed into a sterile flask and shaken in a 0.5% sodium hypochlorite solution for 4 minutes. This mix was then poured over a clean stack of 8” sieves (No. 270 over No. 500) and rinsed to collect any eggs. The collected contents of the No. 500 sieve were used to inoculate the susceptible tomato ‘Rutgers’ planted in autoclave-sterilized soil. Tomatoes were grown under the same greenhouse conditions as described in Section 2.1. Eight weeks post inoculation, the roots of each tomato plant were rinsed and examined for evidence of RKN galling or egg masses. If present, at least three RKN females were individually excised from the plant roots and extracted for DNA analysis as previously described (Holterman et al. 2012).

### 2.6 Diagnostic PCR and Sanger sequencing for individual females

RKN species identification of individual females was conducted using two mitochondrial specific primer sets TRNAH/MRH106 and MORF/MTHIS, and *M. enterolobii* specific primer set MeF/MeR as previously described (Baidoo et al. 2016, Hu et al. 2011, Pagan et al. 2015). We used amplicon sequencing to confirm the species of 2-3 females from the *post-hoc* assays that were found to not be *M. enterolobii*. Using the method described by Janssen et al. (2016), the cytochrome C oxidase subunit II gene was amplified with the Cox2 primer set. PCR reactions were conducted using the Q5® High-Fidelity DNA polymerase system (New England BioLabs, Ipswich, MA); 5 μL 5X Q5 Reaction Buffer,0.25 μL Q5 High Fidelity DNA Polymerase, 200 μM dNTP mix, 1.25 μL each of 10-μM forward and reverse primer for Cox2 and 3 μL DNA template from a single female were combined to a final volume of 25 μL (see Supplemental Tables 1 & 2 for primer sequences and reaction conditions). The sizes of PCR products were first confirmed using gel electrophoresis on a 1.5% agarose gel and then amplicons were sent for Sanger sequencing at Eton Bioscience Inc. (Eton Bioscience Inc., Research Triangle Park, NC) using both forward and reverse primers. Consensus sequences were assembled in Geneious Prime® 2022.1.1 (https://www.geneious.com) and aligned with published mtDNA sequences of tropical RKN available on NCBI GenBank (Supplementary Table 3).

## 3. Results

### 3.1 RKN can be detected in total DNA collected from infected fresh market sweetpotato skins

We collected six individual fresh market sweetpotatoes that displayed symptoms of RKN infection (Fig. 2a). Cross-sections of these infected sweetpotatoes indicated that RKN females were concentrated in the outer ‘skin’ of the roots, from the perimedulla to the periderm (Fig. 2b). We used well established methods to excavate individual *Meloidogyne* females from each root and used diagnostic PCR to identify all females as *M. enterolobii* (Fig. 2c). To determine if these results could be confirmed using a faster approach, we separately skinned each root using a handheld vegetable peeler and extracted total DNA from the skin tissue. We tested these DNA samples using the same diagnostic PCR primers used to speciate individual females (Fig. 2d). The results of each diagnostic PCR confirmed the presence of *M. enterolobii* in each skinning sample from the six roots tested.

**Figure 2.**
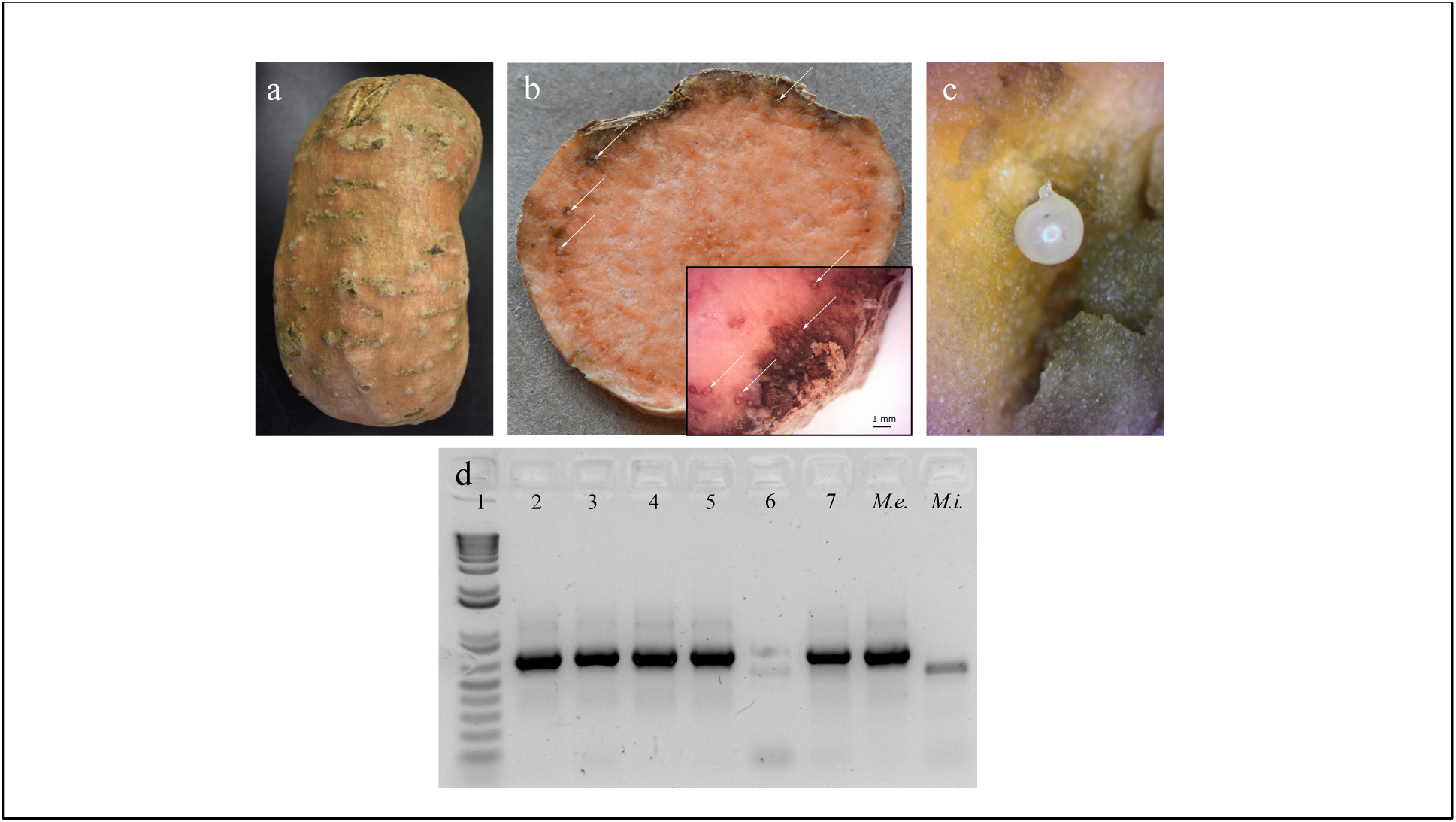
Signs and symptoms of root-knot nematode infection in sweetpotato, whose presence can be detected using a combination of a total skin DNA extraction and PCR. (a) An example of a severely infected sweetpotato storage root collected from a grocery store in Charleston, SC, and was found to have originated from a producer in another state. (b) A cross sectional view of a storage root infected with *M. enterolobii*. Females (white arrows) are concentrated on the outer portions of the root, just under the skin. (c) An example of an *M. enterolobii* female manually excavated from the skin of a sweetpotato. (d) An agarose gel displaying results from a PCR performed on total DNA extracted from six skinning samples collected from individual fresh market sweetpotatoes known to be infected with *M. enterolobii*. PCR was performed with primer set TRNAH/MRH106, which produces a 722bp amplicon from *M. enterolobii* (Lane 8) and a 550bp amplicon from *M. incognita* (Lane 9). Lane 1: 1Kb+ ladder, lanes 2-7: Total skin DNA collected from individual symptomatic sweetpotatoes, Lane 8: Individual *M. enterolobii* female, Lane 9: Individual *M. incognita* female.

### 3.2 Species specific PCR can be used for sensitive detection of *Meloidogyne enterolobii* in skinning samples collected from batches of sweetpotato storage roots

Based on the results from individual roots, we hypothesized that we could scale up this approach to identify *M. enterolobii* in skins collected from multiple sweetpotatoes at once. To test this, we developed a method to quickly remove and collect homogenized skin tissue from batches of sweetpotatoes of up to 10 lbs (Supplemental Film 1). We tested the efficacy of this method using both positive control storage roots (infected with *M. enterolobii* in a greenhouse) and negative control storage roots (grown in sterilized soil in a greenhouse). We used the skin sampling method to process these control roots in 4 different combinations (i.e. control batches) with varying levels of infection: 1) heavily infected control batches (Fig. 3a), 2) moderately infected control batches (Fig. 3b), 3) minimally infected control batches (Fig. 3c), and 4) uninfected control batches (Fig. 3d).

**Figure 3.**
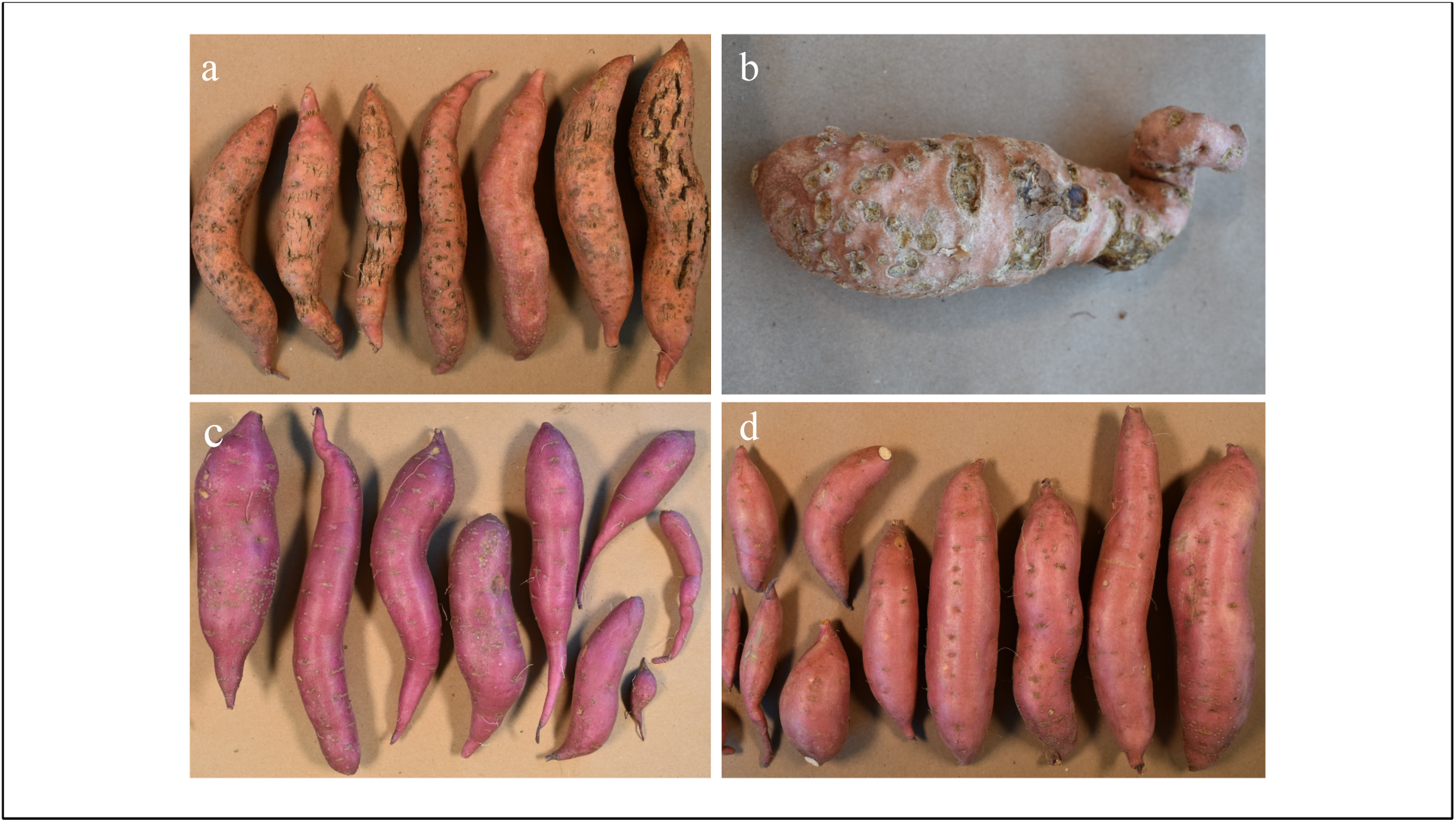
Control storage roots used to test the skinning collection and detection approach; (a) An example of a heavily infected control batch of susceptible (‘Beauregard’) sweetpotatoes inoculated in a greenhouse and used. (b) An example of an individual *M. enterolobii* infected susceptible (‘Beauregard’) root added that was added into a batch of non-symptomatic roots to produce a moderately infected control batch. (c) An example of a minimally infected batch of resistant (‘Regal’) sweetpotatoes which showed little to no galling symptoms used as positive controls. (d) Uninfected (clean) sweetpotatoes used as negative controls.

Using *M. enterolobii* specific qPCR, we detected *M. enterolobii* in all 22 positive control batches regardless of their level of infection (Table 1). Moreover, we observed a negative correlation between the level of batch infection type and the number of qPCR cycles (Ct value) required to detect the presence of *M. enterolobii* DNA. The mean Ct value across all heavily infected batches (24.8) was 1.6 times smaller than the mean Ct value for all minimally infected batches (40.6). Conversely, *M. enterolobii* was not detected in any of the 21 uninfected negative control batches tested with this method (Table 1).

**Table 1.** Control sweetpotato storage root batches used for validating the skin-based survey approach for detecting *Meloidogyne enterolobii*.

| Batch type | Skin-based detection (qPCR) |  | Mean qPCR threshold values<sup>a</sup> | Post-hoc assay detection<sup>b</sup> |  |
| --- | --- | --- | --- | --- | --- |
|  | + | - |  | + | - |
| Uninfected | 0 | 21 | No amplification | 0 | 14 |
| Heavily infected | 6 | 0 | 24.8 ± 1.58 | n/a | n/a |
| Moderately infected | 4 | 0 | 34.3 ± 1.53 | 4 | 0 |
| Minimally infected | 12 | 0 | 40.6 ± 1.77 | 0 | 6 |

<sup>a</sup> Mean Ct value ± standard deviation
<sup>b</sup> A *post-hoc* inoculation assay was conducted by using eggs extracted from the remaining skinnings of each batch to inoculate susceptible tomato plants in the greenhouse.

To confirm the presence or absence as well as the infectivity of *M. enterolobii* collected from these control skinning batches, we performed *post-hoc* infection assays on select control batches. Using the remaining skinnings from each selected control batch we inoculated susceptible tomato plants in the greenhouse. The results of these *post-hoc* assays largely confirmed the qPCR results. *M. enterolobii* infection was found on all four tomato root systems inoculated with skinnings from moderately infected control batches. No infection was found on any of the tomatoes inoculated with the 14 uninfected control batches. Interestingly, no infection was found on any of the six tomato roots inoculated with skinnings from the minimally infected control batches containing *M. enterolobii* resistant sweetpotatoes (’Regal’ and USDA-10-185) (Table 1). These findings indicate the method was sensitive enough to detect the presence of *M. enterolobii* DNA in sweetpotato batches that were both asymptomatic and non-infective.

To further test the sensitivity of the total skin sampling method, we performed qPCR on DNA from inoculated skin samples ranging in concentration from 0 to 100 *M. enterolobii* eggs per 10 mL skinning sample. We again observed a negative correlation between the concentration of nematode eggs within a skin sample and the detection Ct value (R^2^=0.90, Fig 4). No amplification was observed in the 3 uninfected negative control samples (0 eggs/10 mL). *M. enterolobii* DNA was amplified in all 3 biological replicates tested at 100 eggs, 10 eggs, and 2 eggs per 10 mL sample concentrations.

**Figure 4.**
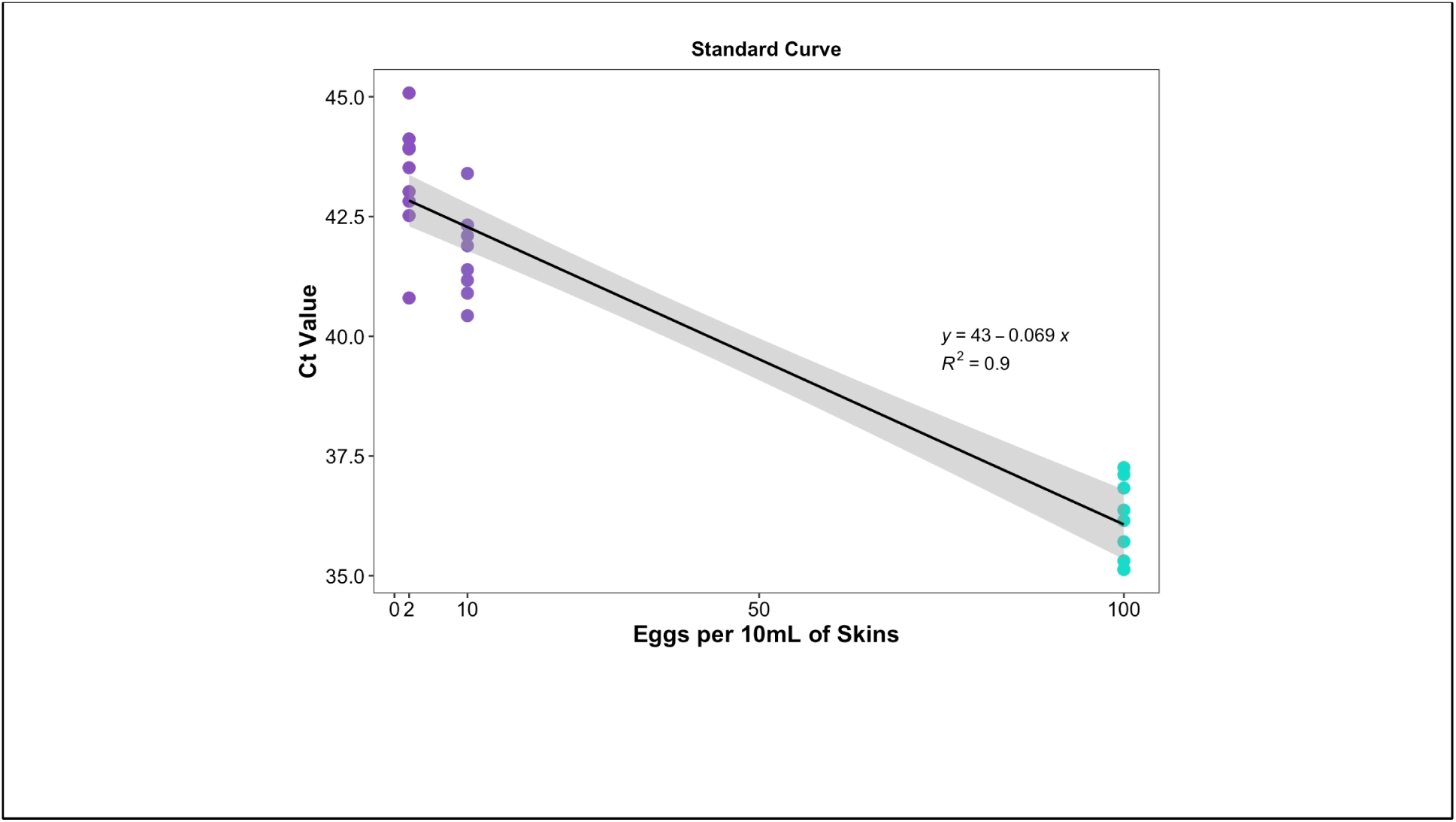
Standard curve generated by amplifying varying concentrations of eggs (0, 2, 10, and 100) per 10 mL of sweetpotato skin samples with *M. enterolobii* specific qPCR primers Ment17F and Ment17R and probe LNA Probe 17 (Kienwick et al. 2015). Each dot represents a technical replicate from one of three biological replicate. The line represents a linear regression model predicting eggs per 10 mL skin sample with the equation y = 43 – 0.069x (R^2^ = 0.9). The shaded region around the line represents a 95% confidence interval.

### 3.3 Skinning based survey of sweetpotato storage roots reliably detected infective RKN in naturally infected roots

To test the efficacy the newly developed skin sampling method to detect RKN in naturally infested sweetpotato samples, we used this method to survey batches of sweetpotatoes collected from both grocery stores and directly from producer fields in South Carolina (Table 2). Of the 30 sweetpotato batches surveyed, RKN were detected in 12 batches and *M. enterolobii* was detected in 3 of those batches.

**Table 2.** Summary of survey of root-knot nematodes using total DNA extracted from sweetpotato storage root skinnings from different sources.

| Source of storage root | Total skin DNA species ID | *Post-hoc* assay species ID<sup>a</sup> | County, State |
| --- | --- | --- | --- |
| Grocery store A | *Meloidogyne* spp. | *M. incognita* | Charleston, SC |
| Grocery store B | *Meloidogyne* spp. | *M. incognita* | Charleston, SC |
| Grocery store C | *M. enterolobii*,<br>*Meloidogyne* spp. | *M. incognita* | Charleston, SC |
| Grocery store D | *Meloidogyne* spp. | *M. incognita* | Sioux Falls, SD |
| Grocery store E | *Meloidogyne* spp. | No infection | Charleston, SC |
| Farmers market A | *Meloidogyne* spp. | No infection | Charleston, SC |
| Farmers market B | *Meloidogyne* spp. | No infection | Charleston, SC |
| Farmers market C | *M. enterolobii*,<br>*Meloidogyne* spp. | *Meloidogyne incognita* | Charleston, SC |
| SC Field A | None | No infection | Sumter, SC |
| SC Field B | None | No infection | Clarendon, SC |
| SC Field C | None | No infection | Clarendon, SC |
| SC Field D | *M. enterolobii* | *M. enterolobii* | Sumter, SC |
| SC Field E | None | No infection | Clarendon, SC |
| SC Field F | None | No infection | Clarendon, SC |
| SC Field G | *Meloidogyne* spp. | No infection | Clarendon, SC |
| SC Field H | *Meloidogyne* spp. | *M. incognita* | Clarendon, SC |
| SC Field I | *Meloidogyne* spp. | *M. incognita* | Clarendon, SC |
| SC Field J | None | No infection | Barnwell, SC |
| SC Field K | None | No infection | Barnwell, SC |
| SC Field L | None | No infection | Barnwell, SC |
| SC Field M | None | No infection | Barnwell, SC |
| SC Field N | None | No infection | Barnwell, SC |
| SC Field O | None | No infection | Barnwell, SC |
| SC Field P | None | No infection | Barnwell, SC |
| SC Field Q | None | No infection | Barnwell, SC |
| SC Field R | None | No infection | Barnwell, SC |
| SC Field S | None | No infection | Barnwell, SC |
| SC Field T | None | No infection | Barnwell, SC |
| SC Field U | None | No infection | Barnwell, SC |

<sup>a</sup> Traditional species identification was used to identify at least three individually extracted females to the species level by amplicon sequencing.

The results of the skinning surveys were confirmed by conducting molecular species identification of females collected from *post-hoc* infection assays (Fig. 5a-b). Identification of individually excised females was possible in 8 out of the 12 batches and was consistent with the original species detection from the skinning sample. For the remaining four batches, there was no evidence of RKN reproduction on the inoculated *post-hoc* root systems. Importantly, we observed no instances of false negative samples in our surveys, i.e., we never found an instance of RKN infection on a *post-hoc* root system that was not detected in the original skin sample.

**Figure 5.**
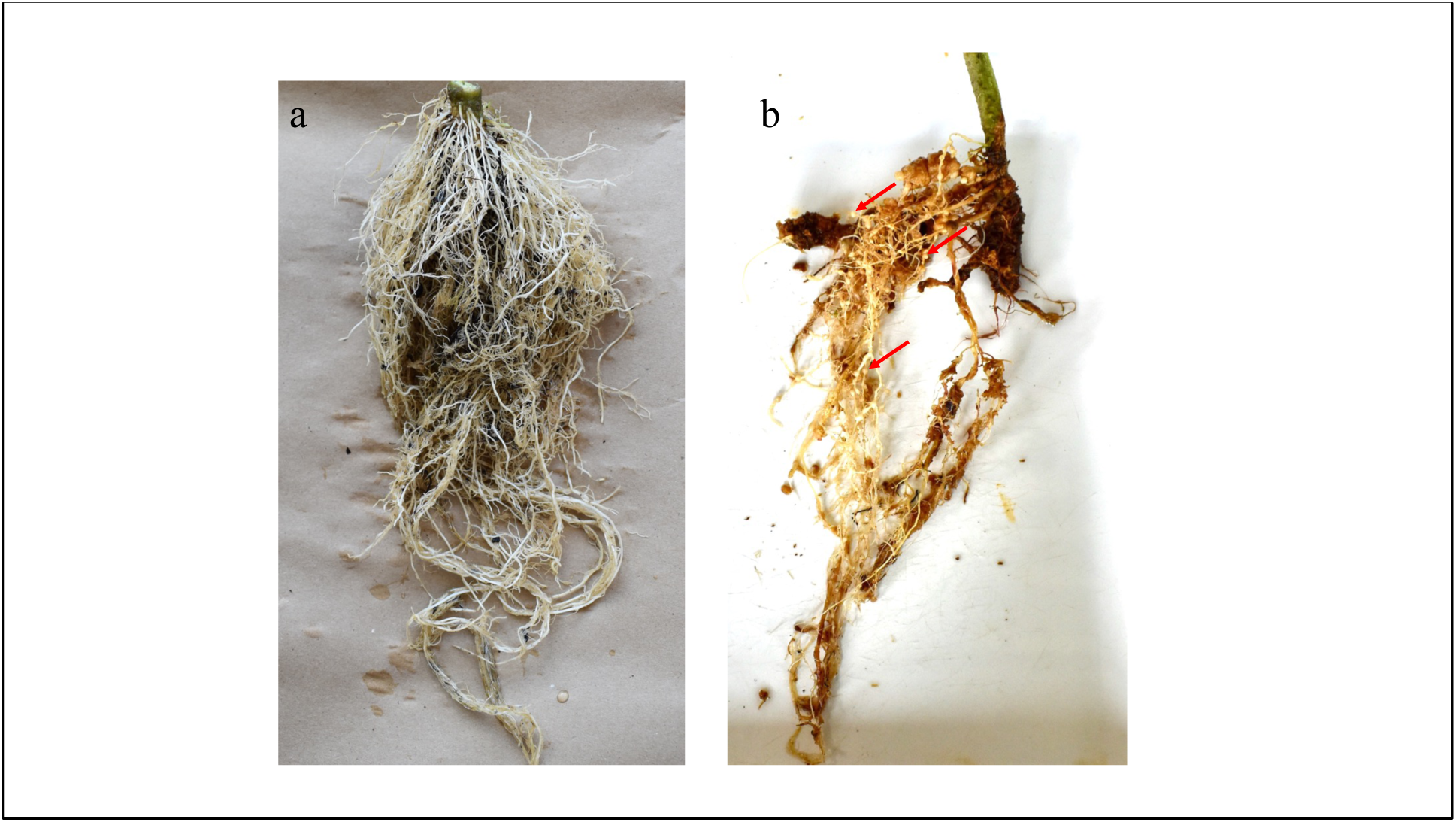
*Post-hoc* bioassay results: roots of susceptible host tomato cultivar Rutgers shown 8 weeks post inoculation with eggs extracted from sweetpotato skinning samples. (a) Tomato roots inoculated with clean control storage root skinning sample, free of any galling. (b) Tomato roots inoculated with a root skinning sample from a moderately infected sweetpotato batch, containing a single heavily *M. enterolobii* infected root showed signs of heavy galling and infection. Red arrows are highlighting individual ssgalls.

## 4. Discussion

We have developed a faster and more efficient method for detecting RKN in batches of sweetpotato storage roots. Using a total skin DNA extraction approach combined with diagnostic PCR provides a quicker and more sensitive alternative to the traditional method of extracting individual nematodes for molecular species identification. In our procedure, the average skinning sample collection time is estimated to be 5-10 min per batch, which is comparable to the amount of time it takes to find, dissect, and remove a single female under a microscope.

In addition to being more time efficient, our results indicate that this total DNA-based detection approach is more sensitive to low levels of infection than traditional methods. Using qPCR, we were able to detect *M. enterolobii* in sweetpotato skin samples which had been inoculated with as little as 2 eggs per 10 mL sample. Additionally, this sampling method allowed for the detection of *M. enterolobii* in resistant host tissue that did not have visible signs of infection and did not produce an infection in our *post-hoc* assays. It would be difficult to find individual nematodes in these resistant roots that could be speciated using traditional methods.

The results of our surveys confirmed the high sensitivity of this screening method, as well as its utility for identifying RKN infection on real world storage root samples. We detected RKN in 12 of 29 sweetpotato batches screened, and we were able to recover RKN in *post-hoc* assays conducted on 8 of these positive samples. The remaining 4 RKN positive sweetpotato batches showed negative results in the *post-hoc* assays. Though these could be thought of as false positives, we suspect that these roots may have been infected with RKN at one point. This scenario would be similar to the negative results we observed in the *post-hoc* assays conducted using our minimally infected control batches, which contained inoculated roots from resistant sweetpotato lines. Moreover, the fact that we never observed a false negative result in any of our other surveys batches confirmed that this is an accurate and reliable method for detecting RKN in sweetpotato storage roots.

We obtained consistent results using species specific primer sets to detect *M. enterolobii* in our skin samples. However, we found that species specific primer sets for other RKN species were less reliable on these same samples. Many of these SCAR primer sets have been designed and tested to speciate individually extracted RKN, but we observed non-specific amplification when they were used on the skinning samples. We speculate that this may be due to the presence of a large number of other microbes that inhabit sweetpotato skins. The total DNA extracted from sweetpotato skins likely contains both sweetpotato DNA as well as that of a plethora of other microbes which could have resulted in non-specific amplification products (Puri et al. 2019). Further research will be needed to design and test other primer sets that can reliably detect other RKN species in these types of metagenomic skin DNA samples.

The community level data collected using this total skin DNA extraction method could help regulators monitor the movement of other soil-borne pathogens on ‘seed’ storage roots. This approach increases the likelihood that a specific species of interest, in this case *M. enterolobii*, will be detected in mixed populations. In fact, we observed several instances of mixed RKN populations within our storage root surveys, where *M. enterolobii* was detected alongside other RKN species (Table 2). It’s possible that other soil-borne pathogens that reside on storage root skins could also be captured using this same sampling method, and other groups have noted the potential utility of metagenomic surveillance approaches for other plant pathogens (Piombo et. al. 2021). Though further study is needed to validate the use of this approach for other sweetpotato pathogens, our results indicate that this is an effective method to screen for *M. enterolobii* infecting storage roots.

*M. enterolobii* has become a serious threat to the production of sweetpotatoes and other vegetables around the world and has quickly spread through the sweetpotato industry in the Southeastern US (FINDMe 2022). It has been proposed that the movement of contaminated plant materials (i.e., seed roots and/or slips) used for propagating sweetpotato could be vectoring *M. enterolobii* between fields, counties, and states (Quesada-Ocampo 2018, 2019). The results of this study support this supposition, showing that viable *M. enterolobii* eggs can be isolated from infected fresh market sweetpotatoes. This further highlights the need for a robust clean seed certification program for sweetpotato to slow the spread of *M. enterolobii* and other soilborne diseases to non-infested sweetpotato fields. This new sweetpotato screening method could be used to help with this seed sweetpotato certification process, by accurately and consistently detecting *M. enterolobii* and other RKN in batches of sweetpotato in a more rapid and cost-effective manner.

## Acknowledgements

We would like to thank the labs lead by Dr. Sebastian Kiewnick and Dr. Janete Brito for their assistance in acquiring and confirming the functionality of *M. enterolobii* specific the qPCR probes used in this study.

