## Supplemental Tables 1-2-3. for "A High-Throughput Sampling Method for Detection of *Meloidogyne enterolobii* and Other Root-Knot Nematodes in Sweetpotato Storage Roots"

**Supplemental Table 1.** Primer sets and LNA probe used for the identification of RKN collected from storage root skinning samples and *post-hoc* tests.

| Primer Name | RKN | Primer sequence 5' – 3' | Fragmen t (bp) | References |
|---|---|---|---|---|
| **TRNAH** | Nonspecific | TGAATTTTTTATTGTGATTAA | 557-720 | (Stanton et al. 1997) |
| **MRH106** | Nonspecific | AATTTCTAAAGACTTTTCTTAGT | | (Stanton et al. 1997) |
| **MORF** | Nonspecific | ATCGGGGTTTAATAATGGG | 743 | (Hugall et al. 1994) |
| **MTHIS** | Nonspecific | AAATTCAATTGAAATTAATAGC | | (Hugall et al. 1994) |
| **MeF** | *M. enterolobii* | AACTTTTGTGAAAGTGCCGCTG | 250 | (Long et al. 2006) |
| **MeR** | *M. enterolobii* | TCAGTTCAGGCAGGATCAACC | | (Long et al. 2006) |
| **Cox2-F** | Nonspecific | TTGAATTTAAGTGTTGTTTATTAC | 432 | (Janssen et al. 2016) |
| **Cox2-R** | Nonspecific | GATTAATACCACAAATCTCTGAAC | | (Janssen et al. 2016) |
| **Ment17F** | *M. enterolobii* | TGT GGT GGC TCA TTT TCA TTA | qPCR | (Kiewnick et al. 2015) |
| **Ment17R** | *M. enterolobii* | AAA AAC CCT AAA AAT ACC CCA AA | qPCR | (Kiewnick et al. 2015) |
| **LNA Probe 17**[a] | *M. enterolobii* | 56-FAM/ A+G+G+A+G+C+T+G /3BHQ_1 | qPCR | (Kiewnick et al. 2015) |

[a]Plus sign located before base pair denotes locked nucleic acid design.

**Supplemental Table 2.** PCR amplification conditions of primers used for molecular identification of RKN.

| Primer Set | Amplification Conditions | |
|---|---|---|
| **TRNAH/MRH106 MORF/MTHIS** | 94 °C 3 min | |
| | 94 °C 30 sec | x 40 cycles |
| | 50 °C 30 sec | |
| | 68 °C 1 min | |
| | 68 °C 10 min | |
| **MeF/MeR** | 94 °C 3 min | |

| | | |
|---|---|---|
| | 94 °C 30 sec | |
| | 64 °C 30 sec | x 35 cycles |
| | 68 °C 30 sec | |
| | 68 °C 10 min | |
| **COX2-F/COX2R** | 98 °C 30 sec | |
| | 98 °C 3 10 sec | |
| | 55 °C 30 sec | x 35 cycles |
| | 72 °C 30 sec | |
| | 72 °C 2 min | |
| **Ment17F/Ment17R LNA Probe 17** | 95 °C 5 min | |
| | 95 °C 10 sec | |
| | 60 °C 1 min | x 45 cycles |
| | 72 °C 3 sec | |
| | 40 °C 10 sec | |

**Supplementary Table 3.** GenBank accession information used for species level identification of individual RKN females used for amplicon sequencing.

| *Meloidogyne spp.* | *GenBank Accession* | *Sequence Origin* | *Primers* | *Reference* |
|---|---|---|---|---|
| *M. incognita* | KJ476151 | Cytochrome C Oxidase Subunit II | COX2F/COX2R | (Janssen et al. 2016) |
| *M. javanica* | KP202352 | Cytochrome C Oxidase Subunit II | COX2F/COX2R | (Janssen et al. 2016) |
